# Prevalence estimate of blood doping in elite track and field at the introduction of the Athlete Biological Passport

**DOI:** 10.1101/736801

**Authors:** Raphael Faiss, Jonas Saugy, Alix Zollinger, Neil Robinson, Frédéric Schütz, Martial Saugy, Pierre-Yves Garnier

## Abstract

In elite sport, the Athlete Biological Passport (ABP) was invented to tackle cheaters by monitoring closely changes in biological parameters, flagging atypical variations. The haematological module of the ABP was indeed adopted in 2011 by the International Association of Athletics Federations (IAAF). This study estimates the prevalence of blood doping based on haematological parameters in a large cohort of track & field athletes measured at two international major events (2011 & 2013 IAAF World Championships) with a hypothesized decrease in prevalence due to the ABP introduction.

A total of 3683 blood samples were collected and analysed from all participating athletes originating from 209 countries. The estimate of doping prevalence was obtained by using a Bayesian network with seven variables, as well as “doping” as a variable mimicking doping with low-doses of recombinant human erythropoietin (rhEPO), to generate reference cumulative distribution functions (CDFs) for the Abnormal Blood Profile Score (ABPS) from the ABP.

Our results from robust haematological parameters indicate an estimation of an overall blood doping prevalence of 18% in average in endurance athletes (95% Confidence Interval (C.I.) 14-22%). A higher prevalence was observed in female athletes (22%, C.I. 16-28%) than in male athletes (15%, C.I. 9-20%). In conclusion, this study presents the first comparison of blood doping prevalence in elite athletes based on biological measurements from major international events that may help scientists and experts to use the ABP in a more efficient and deterrent way.

**What are the new findings ?:**

- This study presents the first comparison of blood doping prevalence in elite track & field athletes based on biological measurements from major international events
- Our results from robust haematological parameters indicate an estimation of an overall blood doping prevalence of 18% in average in endurance athletes.
- The confidence intervals for blood doping prevalence range from 9-28% with wide discrepancies between certain countries.

**How might it impact on clinical practice in the near future:**

- The further development of the Athlete Biological Passport with a careful monitoring of biological parameters still represents the most consistent approach to thwart athletes using undetectable prohibited substances or methods.
- This study describes a method to define blood doping prevalence with the analysis of robust haematological parameters
- Estimates of doping prevalence in subpopulations of athletes may represent a valuable tool for the antidoping authorities to perform a risk assessment in their sport.

## Introduction

The true prevalence of doping among athletes competing at the highest level remains virtually unknown while few attempts to address this point exist (Scarpino et al. 1990, Sottas, Robinson, Fischetto, et al. 2011, Striegel et al. 2010, Thevis et al. 2008, Ulrich et al. 2018). Prevalence of doping in sports is influenced by many cultural, environmental or social factors, and the efficiency of the anti-doping strategy is an important feature influencing this prevalence (Sjoqvist et al. 2008). In fact, official adverse or atypical results occur in less than 2% of the tests performed in laboratories accredited by the World Anti-Doping Agency (WADA) (WADA 2018). However, such statistic is flawed and does not allow an estimate of the prevalence of doping in athletes for at least two reasons. First, because drug tests give priority to specificity rather than sensitivity, false-negative results lead to underestimate the true values because of a lack of sensitivity (Sottas, Robinson, Fischetto, Dolle, Alonso and Saugy 2011). In addition cheats using low dosages of doping substances result in very short detection windows (de Hon et al. 2015). Second, some tests (e.g., in competition or at random) may have a primary deterrent effect rather than being able to detect cheats immediately. Surveys of athletes may represent an attractive alternative while truthful answers from top-level athletes tempted to deflect any suspicions towards themselves or their sport are far from guaranteed. For example, doping (in all forms) prevalence is said to range between 39% and 62% based on anonymous questionnaires answered by athletes competing in two 2011 competitions of the International Association of Athletics Federations (Ulrich, Pope, Cleret, Petroczi, Nepusz, Schaffer, Kanayama, Comstock and Simon 2018). Notwithstanding the surprisingly big values, these results shall first underline the large variability and heterogeneity in the determination of doping prevalence with a significance close to any wet finger approach.

In another way, performance data from athletes convicted for doping violations was used to assess the predictive performance of a Bayesian framework with a probit model. Such a model was able to detect performance differences between doped and presumed clean shotput athletes (Montagna and Hopker 2018). The latter supports the robustness of objective data (e.g., measurable performance or haematological variables) for an unbiased estimate of doping prevalence in sport.

In this context, monitoring an athletes’ haematological parameters is a smart concept allowing to track individual changes overtime with discrepancies naturally due in a certain range to physiological changes and potentially due to any external cause (medical condition or doping) over a certain limit. Such a concept of longitudinal monitoring of blood parameters was conceived in parallel to direct detection methods with a mathematical model to identify biological markers indicative of doping with the Athlete Biological Passport (ABP) (Salamin et al. 2017, Saugy et al. 2014, Striegel, Ulrich and Simon 2010). The ABP is thus based on a longitudinal approach of individual changes in selected biomarkers using a Bayesian statistical method. Nowadays, the high standardization of the blood tests according to ABP guidelines published by the WADA (WADA 2009) would allow a reliable estimation of blood doping prevalence through epidemiological measures of occurrence. For example, the Abnormal Blood Profile Score (ABPS) is calculated and quantified in the ABP. The ABPS is calculated based on a combination of reticulocyte percentage (RET%), red blood cell count (RBC), haemoglobin (HGB), haematocrit (HCT), mean corpuscular volume (MCV), mean corpuscular haemoglobin (MCH), and mean corpuscular haemoglobin concentration (MCHC) (Schütz and Zollinger 2018, Sottas et al. 2007). This score has been successfully used to estimate the prevalence of blood doping in elite track and field athletes (Sottas, Robinson, Rabin, et al. 2011).

Indeed, the IAAF targeted top-level track and field athletes with complete blood testing programs as early as in 2001 (resulting in the latter study) and adopted the ABP in 2011 after it was first introduced in cycling in 2009. For instance, blood tests for all athletes participating in the 2011 IAAF World Championships in Daegu (Republic of Korea) served to build a solid reference basis for haematological, steroidal and endocrines modules in these athletes (Robinson et al. 2012). The IAAF decided to announce this exceptional testing program prior to the event and repeated the program in the 2013 world championships in 2013 (Moscow, Russia) with all athletes participating being tested for blood parameters. A thorough description of these data allowed the recent publication of a worldwide distribution of blood values in elite track and field athletes (Robinson et al. 2019). All blood parameters determined from these samples collected during the events were then introduced in each athlete’s individual haematological module of the Athlete Biological Passport for further analysis.

Consequently, this study aims to analyse the data collected during the 2011 and 2013 event and to present estimates of the prevalence of doping in participating athletes based on the evaluation of specific blood variables determined for the ABP. With the individual longitudinal monitoring of biological variations implemented in 2011 by the IAAF (i.e. the ABP to scrutinize variations in blood parameters), it is hypothesized to observe a decrease in this doping prevalence between 2011 and 2013. Additionally, the study allows an estimate of the prevalence of blood manipulations, not only in the entire population of endurance athletes, but also in sub-groups (countries) participating to these two competitions. This approach provides important information to the antidoping authority to elaborate an appropriate antidoping policy.

## Subjects and Methods

### Sample collection and biological analyses

A total of 3683 blood samples were collected and analysed during the IAAF World Championships: 1808 in Daegu (Republic of Korea, 2011) and 1875 in Moscow (Russia, 2013). All athletes originating from 209 countries were controlled before the competition. In case some athletes were tested more than once during the competition period, only the first record was kept for our analysis. Blood sampling took place between 07:05 and 24:00 o’clock. A detailed description of the athletes included and the resulting samples included in this study are described in detail elsewhere (Robinson, Saugy, Schutz, Faiss, Baume, Giraud and Saugy 2019). Due to the design of the study collecting blood samples in all athletes who competed in two major events, we may consider that the dataset represents the most comprehensive population possible (maximal sample size). The lack of sample size analysis is justified by the anti-doping perspective of our work hypothesis aimed at avoiding false-positives (high specificity), even though this may result in false-negatives (lower sensitivity). As such, we acknowledge that we may be unable to identify some countries with a non-zero prevalence of doping because sufficient power to identify the effect is lacking due to the sample size limited by the design of the study itself. Sample collection procedures, preconditioning, analysis and storage have been thoroughly described for the 2011 event in Daegu (Robinson, Dolle, Garnier and Saugy 2012) and precisely reproduced in 2013 in Moscow in order to allow for a comparative analysis of the results. However, for convenience to the reader, we describe here some key points to set the context of the present study. For the events, a mobile WADA-accredited laboratory unit was created including several blood collection stations. Athletes were requested to report to a designated station within 24 h upon their arrival on the competition sites as part as the routine anti-doping procedure. All blood tests were done following the WADA ABP operating guidelines (WADA 2009) and tubes were stored immediately in monitored fridges before transportation to the on-site analytical laboratory in an insulated cool box with a controlled temperature of approximately 4°C (as recorded by a temperature datalogger). All samples were analysed within less than 24 hours after blood withdrawal with one of the two identical haematological analysers (Sysmex XT-2000i, Sysmex Europe, Norderstedt, Germany) required to manage the high number of samples. In Daegu, the mobile unit was managed by a team from the Lausanne WADA accredited Laboratory whereas in Russia, the analyses were conducted by the Moscow WADA accredited laboratory, in accordance with the related WADA technical document (WADA 2009).

The ABPS was calculated from 7 haematological variables: RET%, HGB, haematocrit (HCT), red blood cell count (RBC), mean corpuscular volume (MCV), mean corpuscular haemoglobin (MCH) and mean corpuscular haemoglobin concentration (MCHC) (Schütz and Zollinger 2018, Sottas et al. 2006).

### Estimation of prevalence and statistical analysis

The ABPS of the population studied was compared with a simulated reference athletic population, based on the population described in (Robinson, Saugy, Schutz, Faiss, Baume, Giraud and Saugy 2019). The reference population is characterized by 7 heterogeneous variables, namely endurance, age, sex, ethnicity (*i.e.* continent), altitude, disease, and instrument used for the analysis.

Athletes were first classified into “endurance” and “non-endurance”; “Endurance” comprised all athletes competing in running or walking events with distances equal or longer than 800 m while “Non-endurance” included all athletes competing in jumps, sprints, throws and combined events, as well as distances shorter than 800 m.

Second, age at sampling collection allowed a classification in the three following categories for all athletes: <=19 years, 19-24 years, >= 25 years, while males and females were separated as such.

Then ethnicity was defined with four categories (Caucasian, Asian, African, Oceanian). Since only information about the country of origin of the athletes was available, proportions were estimated for countries having more than 10 athletes based on the Central Intelligence Agency World Factbook (CIA 2018). While such statistics concern the general population, it still provides a fair estimate of athletes representing those countries and should not be considered as a bias to the results. Countries with less than 10 athletes were grouped by continent with imposed proportions of ethnicity (ordered as Caucasian, Asian, African and Oceanian) per continent as follows: Europe=(1,0,0,0), Asia=(0,1,0,0), Africa=(0,0,1,0), North America=(0.48,0.07,0.45,0), South America=(0.25,0.25,0.25,0.25) and Oceania=(0,0,0,1).

Altitude exposure before the events was considered and differentiated between endurance and non-endurance athletes with an allocation of athletes between categories: <1000 m, 1000-1500 m, 1500-2000 m, and >2000 m. Because no information about prior altitude exposure was available in Daegu, following proportions in the respective categories described above were arbitrarily applied to endurance athletes (0.5,0.2,0.2, and 0.1) and non-endurance athletes (0.96, 0.02, 0.01, 0.01). Since prior altitude exposure data was only partially recorded in Moscow, athletes were allocated to the same categories with the following proportions: endurance (0.5329, 0.1557, 0.1557, 0.1557) and non-endurance (0.96, 0.0133, 0.0133, 0.0133).

All athletes were assumed to be healthy (i.e. not sick at the moment of sample collection) and non-smoking.

To estimate the prevalence of doping in different populations, a Bayesian network was used with the 7 variables described above, as well as “doping” as a variable mimicking doping with low-doses of rhEPO, to generate the simulated reference population, which is then used to generate reference cumulative distribution functions (CDFs, solid curves from Figure 1) for the marker ABPS. The generation of CDFs is well described elsewhere (Sottas, Robinson, Fischetto, Dolle, Alonso and Saugy 2011). Briefly, the value of prevalence is estimated from the cumulative distribution function as the ratio of two areas: 1) the area between the reference curve of no-doping (solid, left) and the ABPS curve (dashed), and 2) the area between the two reference curves (solid, left and right) (see Figure 1).

**Figure 1.**
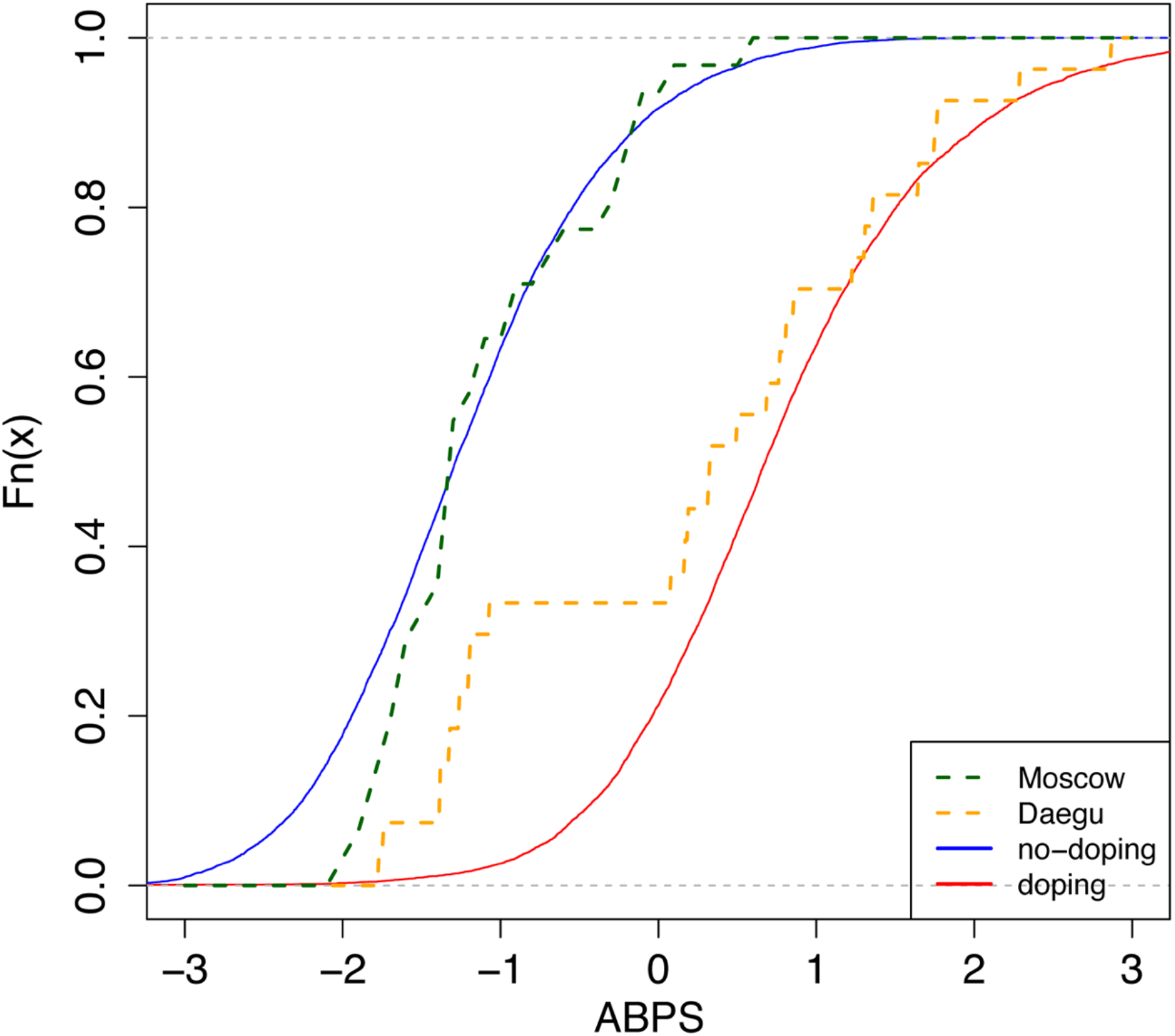
Cumulative distribution functions (CDFs) of the Abnormal Blood Profile Score (ABPS) marker as calculated in the Athlete Biological Passport indicating doping prevalence for endurance athletes from Country N Solid lines: reference CDFs obtained for a modal population of endurance athletes; blue: assuming no doping, red: assuming doping with microdoses of rhEPO (Ma et al. 2008, Sottas et al. 2008, Sottas, Robinson, Fischetto, Dolle, Alonso and Saugy 2011). The difference between both lines refers to the discriminative power of the ABPS marker. Dashed lines: empirical CDFs obtained from all tests performed in endurance athletes in Daegu (orange, n=27) and Moscow (green, n=31)

Besides, in the original paper (Sottas, Robinson, Fischetto, Dolle, Alonso and Saugy 2011), no difference in the variation of the ABPS between men and women was assumed. However, in our study, we observed a striking difference. We therefore introduced a correction factor for sex in the Bayesian network. In addition, reference values were adjusted for the mean and standard deviation (µ, σ of the doping and non-doping reference curves of ABPS) in order to correspond to the observed ABPS score. A systematical shift was observed in all ABPS values measured in Moscow compared to the values measured in Daegu (−0.16); since such a shift is not related to a particular biological reason, it was corrected using the adjustment factor determined from the reference values described in (Robinson, Saugy, Schutz, Faiss, Baume, Giraud and Saugy 2019). Since the seven haematological variables are related, only HGB, HCT, RBC and RET% were corrected for this pre-analytical bias, using the following additive factors: HGB −0.3, HCT −0.7, RBC −0.08, RET% +0.07. The three other variables were calculated as MCV=10xHCT/RBC, MCH=10xHGB/RBC, and MCHC=100xHGB/HCT.

Finally, the time of the day the blood samples were collected induced non-analytical variations in the ABPS. Values of the ABPS for all athletes whose blood samples were taken in the afternoon or in the evening (reference is set to be morning sampling) were additionally corrected. The reference curve corresponds to non-endurance athletes; to ensure that the corrections described above were required and not linked to measurement bias, linear models were used to make sure that the correction factors were the same for endurance and non-endurance athletes (data not shown).

The estimated doping prevalences at Daegu and Moscow were compared using a Kolmogorov-Smirnoff test, which assesses the largest vertical difference between the two curves, and a Cramér von Mises test, which considers the sum of the differences. The p-values obtained were adjusted for multiple testing using the Benjamini-Hochberg procedure to control the False Discovery Rate (Benjamini and Hochberg 1995). The 95% confidence intervals for prevalence were obtained by resampling and constructed based on 1000 bootstrap estimates from the observed data. While the latter analysis is inappropriate in case of zero estimates and may lack sufficient power in the case of small sample sizes, the inter-athlete variability within our population did not produce exact zero estimates and the applied resampling method may thus still provide pertinent results in the particular scenario of our study.

It is worth noting that the use of simulated reference populations (and the absence of an ideal reference population with a known zero prevalence of doping), combined with the inter-variability of measurements across different athletes, can cause some estimates of variability (as well as the extremities of some of the confidence intervals) to be negative. We acknowledge that negative prevalence estimates are impossible to achieve. However, if a population has a 0 % estimate of doping prevalence, ABPS values from the subset population below the median value would produce a negative estimate. The other half of the population would in turn have a positive estimate of prevalence, thus explaining the null prevalence for the entire population.

All calculations and analyses were performed using the R software (R Development Core Team 2005). Values for the Abnormal Blood Profile Score (ABPS) were calculated using a custom-made R package described elsewhere (Schütz and Zollinger 2018).

## Results

### Description of the population analysed

Table 1 presents the distribution of samples across sex, age, continent, sport and altitude variables for the IAAF World Championships in Daegu (2011) and Moscow (2013). Each variable is divided in subcategories. NCC America stands for North America, Central America and Caribbean. A total of 3683 samples were tested, among those the proportions of males versus females were almost equal. In addition, approximately one third were competing in endurance disciplines.

**Table 1.** Proportions of samples for each variable for Daegu (2011) and Moscow (2013) IAAF World Championships. NCC America stands for North America, Central American and Caribbean.

| Variable | Categories | Daegu (n=1808) | Moscow (n=1875) |
| --- | --- | --- | --- |
| Sex | Male | 53% | 56% |
|  | Female | 47% | 44% |
| Age | ≤ 19 | 2% | 2% |
|  | 19-24 | 38% | 40% |
|  | ≥ 25 | 60% | 58% |
| Continent | Africa | 15% | 14% |
|  | Asia | 16% | 12% |
|  | Europe | 42% | 44% |
|  | NCC America | 19% | 20% |
|  | Oceania | 4% | 4% |
|  | South America | 4% | 5% |
| Sport | endurance | 31% | 35% |
|  | non-endurance | 69% | 65% |
| Altitude | < 1000 m | NA | 81% |
|  | > 1000 m | NA | 19% |

NA, no adequate data available

### Prevalence of doping: endurance only

Table 2 presents the prevalence of blood doping along with 95% confidence intervals in all endurance athletes for Daegu and Moscow (569 athletes in Daegu and 653 athletes in Moscow), stratified by sex or country. Only the 18 countries with at least 10 athletes competing in endurance sports at either Daegu or Moscow competitions are represented. The overall prevalence of blood doping does not decrease significantly between Daegu and Moscow (0.18 to 0.15, NS). The overall prevalence of doping between the two competitions indicates a non-significant decrease in female athletes (0.22 to 0.12, NS) while a non-significant increase is observed in male athletes (0.15 to 0.17, NS). Among the selected countries, 8 tend to decrease their prevalence of blood doping between 2011 and 2013 WCS (only significantly for countries N and Q, P<0.001), 8 increase (significantly only for country L) and two of them stay at the same level (0.00 for countries H and I in both competitions).

**Table 2.** Prevalence of blood doping along with 95 % confidence intervals. The prevalences for the countries are given for endurance athletes only. The ‘n’ columns give the number of athletes for each category for Daegu and Moscow competitions. Only countries with more than 10 athletes competing in endurance sports at either Moscow or Daegu competitions are presented

| Sex & country | 2011 Daegu |  |  | 2013 Moscow |  |  | p.adj ks | p.adj cvm |
| --- | --- | --- | --- | --- | --- | --- | --- | --- |
|  | n | prevalence | CI | n | prevalence | CI |  |  |
| All | 569 | 0.18 | [0.14,0.22] | 653 | 0.15 | [0.12,0.19] | 0.80 | 0.31 |
| Female | 246 | 0.22 | [0.16,0.28] | 276 | 0.12 | [0.09,0.16] | 0.80 | 0.31 |
| Male | 323 | 0.15 | [0.09,0.2] | 377 | 0.17 | [0.12,0.22] | 0.80 | 0.31 |
| Country A | 15 | 0.00 | [-0.22,0.09] | 22 | 0.04 | [-0.12,0.2] | 0.98 | 0.50 |
| Country B | 11 | 0.15 | [-0.16,0.45] | 8 | 0.02 | [-0.24,0.27] | 0.98 | 0.48 |
| Country C | 6 | 0.32 | [0.02,0.61] | 18 | 0.07 | [-0.11,0.25] | 0.93 | 0.31 |
| Country D | 8 | 0.17 | [-0.04,0.39] | 10 | 0.31 | [0.06,0.56] | 0.99 | 0.63 |
| Country E | 33 | 0.19 | [0.09,0.3] | 37 | 0.30 | [0.17,0.43] | 0.98 | 0.50 |
| Country F | 10 | 0.03 | [-0.2,0.28] | 14 | 0.17 | [0,0.34] | 0.98 | 0.53 |
| Country G | 16 | 0.04 | [-0.1,0.2] | 14 | 0.18 | [-0.04,0.42] | 0.98 | 0.30 |
| Country H | 7 | 0.00 | [-0.47,0.02] | 18 | 0.00 | [-0.14,0.14] | 0.99 | 0.63 |
| Country I | 25 | 0.00 | [-0.22,0] | 20 | 0.00 | [-0.27,0.2] | 0.98 | 0.31 |
| Country J | 43 | 0.19 | [0.06,0.32] | 42 | 0.13 | [0,0.26] | 0.99 | 0.63 |
| Country K | 17 | 0.47 | [0.17,0.8] | 19 | 0.61 | [0.34,0.86] | 0.98 | 0.53 |
| Country L | 16 | 0.00 | [-0.22,0.09] | 18 | 0.27 | [0.07,0.48] | 0.80 | 0.04 |
| Country M | 27 | 0.15 | [0.01,0.28] | 23 | 0.09 | [-0.06,0.23] | 0.99 | 0.61 |
| Country N | 27 | 0.74 | [0.48,0.99] | 31 | 0.09 | [-0.03,0.2] | <0.001 | <0.001 |
| Country O | 10 | 0.66 | [0.18,1.14] | 6 | 0.23 | [-0.12,0.58] | 0.98 | 0.31 |
| Country P | 10 | 0.17 | [-0.13,0.45] | 11 | 0.11 | [-0.11,0.31] | 0.98 | 0.53 |
| Country Q | 19 | 0.89 | [0.63,1.14] | 16 | 0.25 | [0.03,0.48] | 0.80 | 0.01 |
| Country R | 41 | 0.14 | [0.03,0.25] | 41 | 0.17 | [0.06,0.28] | 0.99 | 0.57 |

CI, Confidence Interval; p.adj ks, p value for the adjusted Kolmogorov-Smirnov test; p.adj cvm, p value for the Cramér–von Mises criterion test

### Prevalence of doping: endurance only for female athletes

Table 3 presents the prevalence of blood doping along with 95% confidence intervals in female athletes for Daegu and Moscow. For pertinence and anonymity, only countries with at least 7 athletes or more competing in endurance sports at either Daegu or Moscow competitions are represented. As said before, the overall prevalence of doping for endurance women only tend to decrease between the two competitions (0.22 to 0.12 for Daegu and Moscow respectively, NS). The prevalence for selected countries only also tends to decrease from 0.30 to 0.15 (P = 0.06). Among selected countries, only four show a decrease in the prevalence of blood doping between 2011 and 2013 (only significant for countries N and Q, P<0.001), 5 tend to increase (not significantly), one stays at the same level (Country I) and 3 of them don’t have sufficient participants to make a prevalence calculation in one of the competitions.

**Table 3.** Prevalence of blood doping along with 95 % confidence intervals for endurance female athletes only. The ‘n’ columns give the number of athletes for each category for Daegu and Moscow competitions. Only countries with more than 7 athletes competing in endurance sports at either Moscow or Daegu competitions are presented.

|  | **2011**<br>**Daegu** |  |  | **2013**<br>**Moscow** |  |  | p.adj<br>ks | p.adj<br>cvm |
| --- | --- | --- | --- | --- | --- | --- | --- | --- |
|  | n | prevalence | CI | n | prevalence | CI |  |  |
| Female endurance | 246 | 0.22 | [0.16,0.28] | 276 | 0.12 | [0.09,0.16] | 0.83 | 0.27 |
| Country A | 4 |  |  | 11 | 0.00 | [-0.11,0.05] |  |  |
| Country C | 2 |  |  | 7 | 0.05 | [-0.11,0.19] |  |  |
| Country E | 16 | 0.21 | [0.1,0.33] | 19 | 0.22 | [0.15,0.3] | 0.93 | 0.70 |
| Country G | 10 | 0.00 | [-0.04,0.04] | 8 | 0.08 | [0.02,0.14] | 0.83 | 0.22 |
| Country I | 13 | 0.00 | [-0.1,0.05] | 7 | 0.00 | [-0.19,0.12] | 0.93 | 0.54 |
| Country J | 21 | 0.10 | [-0.04,0.24] | 19 | 0.09 | [-0.05,0.23] | 0.93 | 0.70 |
| Country K | 6 | 0.41 | [-0.23,1.03] | 7 | 0.24 | [-0.08,0.56] | 0.93 | 0.70 |
| Country L | 9 | 0.00 | [-0.15,0.06] | 10 | 0.19 | [0.01,0.39] | 0.83 | 0.22 |
| Country M | 8 | 0.17 | [0.02,0.33] | 7 | 0.30 | [0.11,0.47] | 0.83 | 0.22 |
| Country N | 15 | 0.73 | [0.43,1.02] | 20 | 0.08 | [-0.04,0.2] | 0.03 | <0.001 |
| Country O | 8 | 0.63 | [0.12,1.14] | 3 |  |  |  |  |
| Country Q | 16 | 0.91 | [0.62,1.21] | 7 | 0.31 | [-0.06,0.66] | 0.21 | <0.001 |
| Country R | 21 | 0.13 | [0.01,0.24] | 22 | 0.19 | [0.11,0.27] | 0.83 | 0.22 |

CI, Confidence Interval; p.adj ks, p value for the adjusted Kolmogorov-Smirnov test; p.adj cvm, p value for the Cramér-von Mises criterion test

### Prevalence of doping: endurance only for male athletes

Table 4 presents the prevalence of blood doping along with 95% confidence intervals in male athletes for Daegu and Moscow. Again, for pertinence and anonymity, only countries with at least 10 athletes or more competing in endurance sports at either Daegu or Moscow competitions are represented. The overall prevalence of blood doping for men in endurance tends to rise between Daegu and Moscow (0.15 versus 0.17, NS). Among selected countries 4 tend to decrease their prevalence of blood doping between 2011 and 2013 (only significantly for Country N), 6 tend to increase (only significantly for Country L), one stays at the same level (Country I, prevalence 0.00) and 2 of them don’t have sufficient participants to make a prevalence calculation in one of the competitions.

**Table 4.**
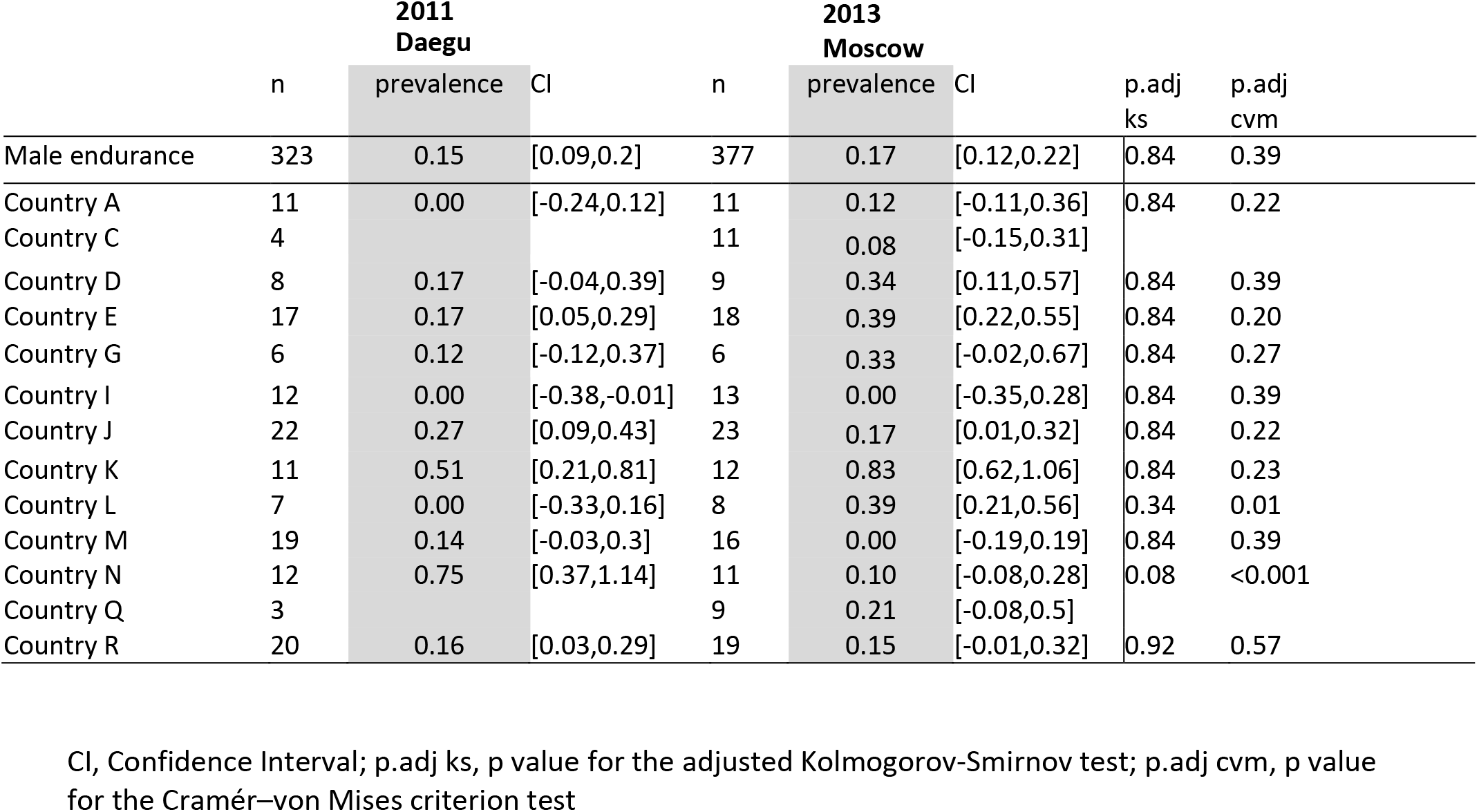
Prevalence of blood doping along with 95 % confidence intervals for endurance male athletes only. The ‘n’ columns give the number of athletes for each category for Daegu and Moscow competitions. Only countries with more than 6 athletes competing in endurance sports at either Moscow or Daegu competitions are presented.

|  | **2011**<br>**Daegu** |  |  | **2013**<br>**Moscow** |  |  | p.adj<br>ks | p.adj<br>cvm |
| --- | --- | --- | --- | --- | --- | --- | --- | --- |
|  | n | prevalence | CI | n | prevalence | CI |  |  |
| Male endurance | 323 | 0.15 | [0.09,0.2] | 377 | 0.17 | [0.12,0.22] | 0.84 | 0.39 |
| Country A | 11 | 0.00 | [-0.24,0.12] | 11 | 0.12 | [-0.11,0.36] | 0.84 | 0.22 |
| Country C | 4 |  |  | 11 | 0.08 | [-0.15,0.31] |  |  |
| Country D | 8 | 0.17 | [-0.04,0.39] | 9 | 0.34 | [0.11,0.57] | 0.84 | 0.39 |
| Country E | 17 | 0.17 | [0.05,0.29] | 18 | 0.39 | [0.22,0.55] | 0.84 | 0.20 |
| Country G | 6 | 0.12 | [-0.12,0.37] | 6 | 0.33 | [-0.02,0.67] | 0.84 | 0.27 |
| Country I | 12 | 0.00 | [-0.38,-0.01] | 13 | 0.00 | [-0.35,0.28] | 0.84 | 0.39 |
| Country J | 22 | 0.27 | [0.09,0.43] | 23 | 0.17 | [0.01,0.32] | 0.84 | 0.22 |
| Country K | 11 | 0.51 | [0.21,0.81] | 12 | 0.83 | [0.62,1.06] | 0.84 | 0.23 |
| Country L | 7 | 0.00 | [-0.33,0.16] | 8 | 0.39 | [0.21,0.56] | 0.34 | 0.01 |
| Country M | 19 | 0.14 | [-0.03,0.3] | 16 | 0.00 | [-0.19,0.19] | 0.84 | 0.39 |
| Country N | 12 | 0.75 | [0.37,1.14] | 11 | 0.10 | [-0.08,0.28] | 0.08 | <0.001 |
| Country Q | 3 |  |  | 9 | 0.21 | [-0.08,0.5] |  |  |
| Country R | 20 | 0.16 | [0.03,0.29] | 19 | 0.15 | [-0.01,0.32] | 0.92 | 0.57 |

CI, Confidence Interval; p.adj ks, p value for the adjusted Kolmogorov-Smirnov test; p.adj cvm, p value for the Cramér–von Mises criterion test

### Comparison of prevalence between Daegu and Moscow

Overall, Figure 2 presents the comparison of blood doping prevalence between Daegu (2011) and Moscow (2013). Figure 2A illustrates prevalence from selected countries without any sex difference, Figure 2B in endurance female athletes only, and Figure 2C in endurance male athletes only.

**Figure 2.**
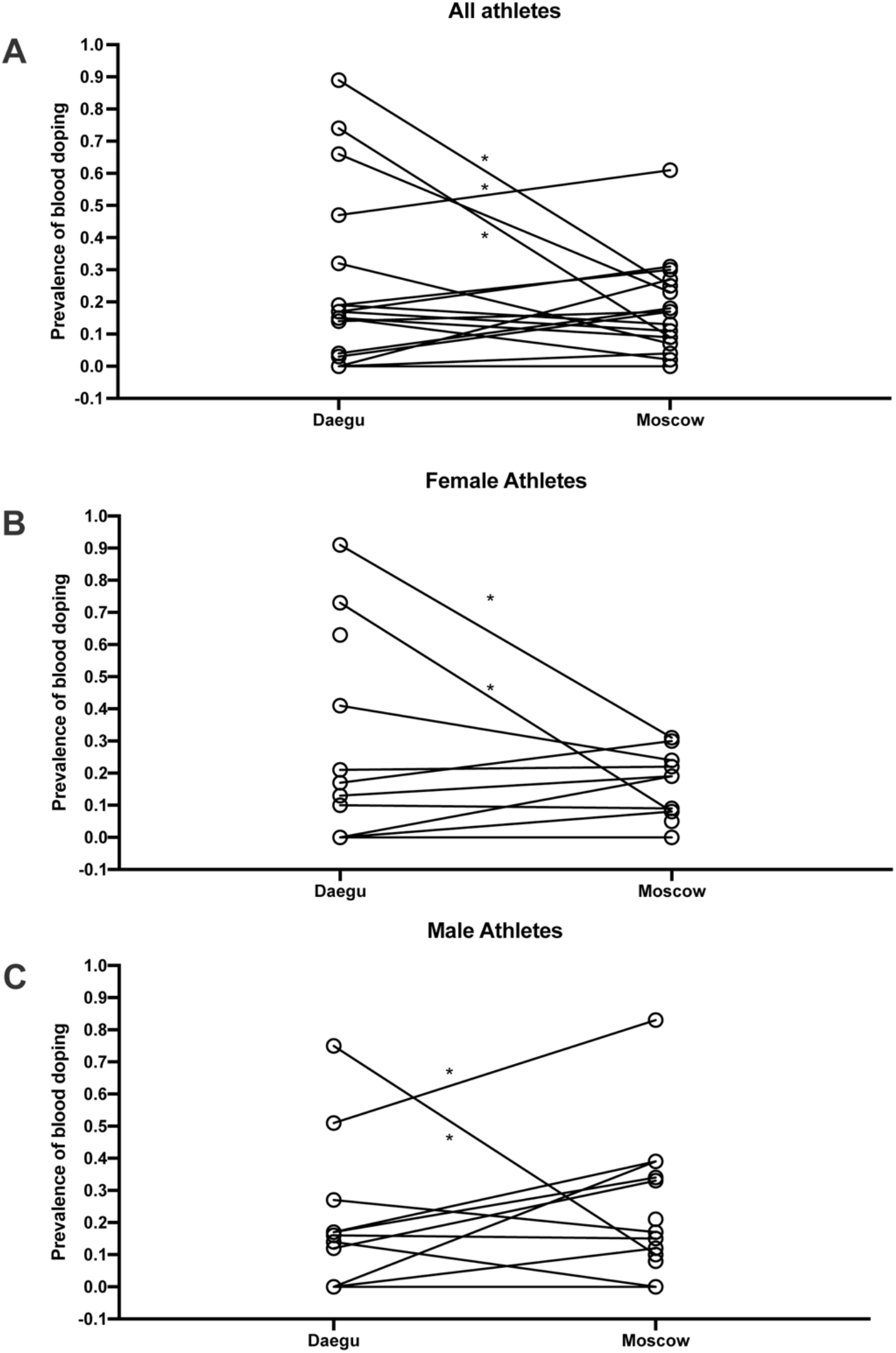
Comparison of blood doping prevalence between Daegu (2011) and Moscow (2013) from selected countries without any sex difference (A), in endurance female athletes only (B), and in endurance male athletes only (C). * P<0.05 for the difference with Daegu

## Discussion

This study presents for the first time a comparison of the prevalence of doping based on blood samples analysed in all athletes competing in two top-level athletic held in 2011 and 2013. The overall estimate of blood doping prevalence in all athletes measured in the 2011 IAAF World Championships indicates that a clear majority of the athletes do not resort to blood doping (i.e., with 18% overall prevalence of doping). This result calculated from accurately determined biological parameters is contrasting with the 44% prevalence for doping in general (i.e. not specifically blood doping) obtained from the same event with a survey-based estimation (Ulrich, Pope, Cleret, Petroczi, Nepusz, Schaffer, Kanayama, Comstock and Simon 2018). In fact, such discrepancy may be first partly explained by the fact that our estimation refers mostly to the prevalence of blood doping that may not be used by non-endurance athletes. Then, the definition of doping differs between both studies. Here we focus specifically on blood doping while Ulrich *et al.* assess doping in general. Indeed, one may understand the clear difference between prevalence calculated from robust biological parameters or self-reported surveys. On one hand, biological variations of haematologic parameters were observed from very strict, controlled and repeatable measurements (thus eliminating acquisition biases). On the other hand, confusion between medication misuse and real doping behaviour for athletes originating from 200 different countries might not be excluded with the analysis of surveys. Blood doping is defined by the WADA as the misuse of certain techniques and/or substances to increase one’s red blood cell mass. An augmented haemoglobin mass is indeed associated with enhanced aerobic performance (Hauser et al. 2016, Saugy et al. 2016) with improved oxygen transport increasing endurance and stamina. The use of an injected Erythropoietin Stimulating Agent (ESA) and blood transfusion (Salamin et al. 2016) may represent the most widespread strategies of blood doping (Elliott 2008, Macdougall and Ashenden 2009). While methods exist to detect ESAs both in blood and urine, autologous blood transfusion remains a challenge in the fight against doping (Morkeberg 2012). While the discovery of erythropoietin (EPO) suddenly made blood doping simpler, many cheaters returned to blood transfusions upon significant development of detection methods (Salamin, Kuuranne, Saugy and Leuenberger 2017, Schumacher et al. 2012). Definitely, autologous blood transfusion is the method of choice with no valid method to date to accurately directly detect such intervention (Malm et al. 2016). Monitoring variations in blood parameters in athletes as implemented with the ABP may however be observed and would allow to flag abnormal alterations. It was therefore hypothesized that the introduction of the ABP by the IAAF in 2011 would result in a decrease of doping prevalence evaluated at the 2013 World championships in the third season after its introduction. Our results indicate that only two countries significantly decreased doping prevalence between 2011 and 2013, one country has a significant increase of prevalence, while the results for the other countries do not support significant changes in the blood doping prevalence. Since all athletes and tests were conducted during the same competitions and under the same protocol, any confounding factors related to procedures or analysis may be excluded. Considering Figure 1, country N produced an empirical CDF (ECDF) very close to the reference CDF representing the “doping” curve in Daegu while its ECDF during Moscow is then very similar to the reference CDF representing the no-doping case. Thus, only external effects may explain the difference between these two ECDFs because the collection, storage and testing protocols were scrupulously the same within the two competitions.

Moreover, since athletes use doping to improve athletic performance and would in turn benefit from a competitive advantage, countries with higher prevalence may obtain better results (e.g., number of medals). For instance, one country improved its ranking among the best countries in the overall medals table despite a significant decrease in doping prevalence observed. Conversely, another country with a low prevalence in Daegu won less medals in Moscow despite a significantly higher doping prevalence. The latter underlines the loophole of prevalence estimates to evaluate competitive results of track and field athletes.

### Limitations and strengths of the study

Our analysis is limited to discrete “in-competition” time-points for the analysis and may not ideally highlight the use of the numerous doping substances or procedures available. While the investigation of doping prevalence is consequently challenging, the use of Bayesian network as used in our study may however yield sufficient power over time to discriminate individual haematological variations (Sottas, Robinson, Rabin and Saugy 2011). In addition, a limitation of this analysis is that the ABPS may not allow to identify all possible blood doping strategies and our results shall thus be interpreted with care. Besides, huge differences between countries are observed with countries where doping prevalence is close to inexistent. The latter is contrasting with WADA statistics underlining adverse analytical findings also in those countries (WADA 2018). This may however be due to stricter anti-doping policies with intelligent and efficient targeting and testing of athletes where the few athletes tempted to use doping are being caught. The number of athletes competing, which is different from one country to another, also influences our ability to assess the prevalence of doping for each country. For countries with a smaller number of athletes competing, the power for detecting differences in doping prevalence will be lower. Given that our dataset includes all athletes competing in two major track & field competitions, the sample size could not be extended. For instance, the anti-doping perspective of our approach aimed at avoiding false-positives (high specificity), even though this may result in false-negatives (lower sensitivity). As such, we acknowledge that the analysis may not identify some countries with a non-zero prevalence of doping because sufficient power to identify the effect is lacking due to the sample size limited by the design of the study itself. Moreover, one should note that the method used to determine prevalence estimates may produce a notable upward bias, because of an unavoidable sparse-data bias due to limited sample size for some countries (Greenland et al. 2016). The results for these countries shall be interpreted with care also considering the descriptive character of this investigation.

In addition, altitude exposure prior to the competitions as a confounding factor was included with due care in the analysis because of its putative influence on the interpretation of ABP data (Lobigs et al. 2018, Sutehall et al. 2019). While data about prior altitude exposure were missing for Daegu and only partially recorded in Moscow, the reference population was constructed with an allocation of 50% of the athletes to some form of prior altitude exposure. Definitely, including more accurate information on prior altitude exposure would help improve the determination of an estimate of blood doping prevalence with the proposed method.

On the other hand, one strength of this study relies in the estimate of prevalence of blood doping based on objective biological parameters. The latter represents a unique opportunity to assess abnormal variations towards a set reference population resulting in a prevalence estimate certainly with lesser bias than survey-based estimates. However, one shall acknowledge that the strength of the analysis relying on a large cohort is being limited by ethnical, sex and environmental factors leading to a lesser power for calculations made comparing unique countries or smaller subgroups of the cohort.

In conclusion, for the first time, an International Federation chose to lead a massive blood testing campaign and announced it. Blood samples from all athletes were analysed in both Daegu (2011) and Moscow (2013) IAAF world championships. Our investigation indicates a moderate prevalence of blood doping in these athletes ranging from 12% to 22%. With the introduction of the ABP in 2011, a decrease in the prevalence of doping over time was hypothesized. However, our results do not support this hypothesis with a tendency for the decrease in doping prevalence (overall from 22% to 12%) between 2011 and 2013. The further development of the Athlete Biological Passport with a careful monitoring of biological parameters still represents the most consistent approach to thwart athletes using undetectable prohibited substances or methods. Such approach may definitely be useful to estimate doping prevalence in subpopulations of athletes, providing a tool for the antidoping authorities to perform a risk assessment in their sport.

## Acknowledgements

The authors wish to acknowledge WADA’s Science Department for the financial support of this study and to the IAAF medical and anti-doping department in Monaco, especially Thomas Capdevielle for the logistic organization of the study. Special recognition is also due to the personal of the Swiss Laboratory for Doping Analyses (LAD) for their technical support.

## Author Contributions Statement

MS, NR and PYG conceived the project and obtained the project funding. MS and NR contributed to the collection of data. AZ and FS statistically analyzed the data. RF, JS and MS drafted the final version of the manuscript. All authors contributed to revising the manuscript and expressed their approval of the final submitted version.

## References

Benjamini Y, Hochberg Y. 1995. Controlling the False Discovery Rate: A Practical and Powerful Approach to Multiple Testing. Journal of the Royal Statistical Society Series B (Methodological). 57:289–300.

The World Factbook. Available from https://www.cia.gov/library/publications/the-world-factbook/fields/2075.html

de Hon O, Kuipers H, van Bottenburg M. 2015. Prevalence of doping use in elite sports: a review of numbers and methods. Sports Med. Jan;45:57–69.

Elliott S. 2008. Erythropoiesis-stimulating agents and other methods to enhance oxygen transport. British journal of pharmacology. Jun;154:529–541.

Greenland S, Mansournia MA, Altman DG. 2016. Sparse data bias: a problem hiding in plain sight. BMJ. 2016-04-27 10:06:06;352.

Hauser A, Schmitt L, Troesch S, Saugy JJ, Cejuela-Anta R, Faiss R, Robinson N, Wehrlin JP, Millet GP. 2016. Similar Hemoglobin Mass Response in Hypobaric and Normobaric Hypoxia in Athletes. Med Sci Sports Exerc. Apr;48:734–741.

Lobigs LM, Garvican-Lewis LA, Vuong VL, Tee N, Gore CJ, Peeling P, Dawson B, Schumacher YO. 2018. Validation of a blood marker for plasma volume in endurance athletes during a live-high train-low altitude training camp. Drug Test Anal. Feb 19. Epub 2018/02/20.

Ma X, Vanasse G, Cartmel B, Wang Y, Selinger HA. 2008. Prevalence of polycythemia vera and essential thrombocythemia. Am J Hematol. May;83:359–362. Epub 2008/01/09.

Macdougall IC, Ashenden M. 2009. Current and upcoming erythropoiesis-stimulating agents, iron products, and other novel anemia medications. Adv Chronic Kidney Dis. Mar;16:117–130.

Malm CB, Khoo NS, Granlund I, Lindstedt E, Hult A. 2016. Autologous Doping with Cryopreserved Red Blood Cells - Effects on Physical Performance and Detection by Multivariate Statistics. PloS one. Jun 10;11.

Montagna S, Hopker J. 2018. A Bayesian Approach for the Use of Athlete Performance Data Within Anti-doping. Frontiers in physiology. 2018-July-19;9.

Morkeberg J. 2012. Detection of autologous blood transfusions in athletes: a historical perspective. Transfus Med Rev. Jul;26:199–208.

R: A language and environment for statistical computing. R Foundation for Statistical Computing [2005. Vienna, Austria.

Robinson N, Dolle G, Garnier PY, Saugy M. 2012. 2011 lAAF World Championships in Daegu: blood tests for all athletes in the framework of the Athlete Biological Passport. Bioanalysis. Jul;4:1633–1643.

Robinson N, Saugy J, Schutz F, Faiss R, Baume N, Giraud S, Saugy M. 2019. Worldwide distribution of blood values in elite track and field athletes: Biomarkers of altered erythropoiesis. Drug Test Anal. Apr;11:567–577. Epub 2018/10/23.

Salamin O, De Angelis S, Tissot JD, Saugy M, Leuenberger N. 2016. Autologous Blood Transfusion in Sports: Emerging Biomarkers. Transfus Med Rev. Jul;30:109–115.

Salamin O, Kuuranne T, Saugy M, Leuenberger N. 2017. Erythropoietin as a performance-enhancing drug: Its mechanistic basis, detection, and potential adverse effects. Mol Cell Endocrinol. Jan 22.

Saugy JJ, Schmitt L, Hauser A, Constantin G, Cejuela R, Faiss R, Wehrlin JP, Rosset J, Robinson N, Millet GP. 2016. Same Performance Changes after Live High-Train Low in Normobaric vs. Hypobaric Hypoxia. Frontiers in physiology.7:138.

Saugy M, Lundby C, Robinson N. 2014. Monitoring of biological markers indicative of doping: the athlete biological passport. Br J Sports Med. May;48:827–U127.

Scarpino V, Arrigo A, Benzi G, Garattini S, La Vecchia C, Bernardi LR, Silvestrini G, Tuccimei G. 1990. Evaluation of prevalence of “doping” among Italian athletes. Lancet. Oct 27;336:1048–1050.

Schumacher YO, Saugy M, Pottgiesser T, Robinson N. 2012. Detection of EPO doping and blood doping: the haematological module of the Athlete Biological Passport. Drug Test Anal. Nov;4:846–853.

Schütz F, Zollinger A. 2018. ABPS: An R Package for Calculating the Abnormal Blood Profile Score. Frontiers in physiology. 2018-November-21;9.

Sjoqvist F, Garle M, Rane A. 2008. Use of doping agents, particularly anabolic steroids, in sports and society. Lancet. May 31;371:1872–1882.

Sottas P-E, Robinson N, Saugy M, Niggli O. 2008. A forensic approach to the interpretation of blood doping markers. Law, Probability and Risk.7:191–210.

Sottas PE, Baume N, Saudan C, Schweizer C, Kamber M, Saugy M. 2007. Bayesian detection of abnormal values in longitudinal biomarkers with an application to T/E ratio. Biostatistics. Apr;8:285–296.

Sottas PE, Robinson N, Fischetto G, Dolle G, Alonso JM, Saugy M. 2011. Prevalence of blood doping in samples collected from elite track and field athletes. Clin Chem. May;57:762–769.

Sottas PE, Robinson N, Giraud S, Taroni F, Kamber M, Saugy M. 2006. Statistical classification of abnormal blood profiles in athletes. Int J Biostat.2:1557–4679. Epub 2006-02-20.

Sottas PE, Robinson N, Rabin O, Saugy M. 2011. The athlete biological passport. Clin Chem. Jul;57:969–976.

Striegel H, Ulrich R, Simon P. 2010. Randomized response estimates for doping and illicit drug use in elite athletes. Drug Alcohol Depend. Jan 15;106:230–232.

Sutehall S, Muniz-Pardos B, Lima G, Wang G, Malinsky FR, Bosch A, Zelenkova I, Tanisawa K, Pigozzi F, Borrione P, et al. 2019. Altitude Training and Recombinant Human Erythropoietin: Considerations for Doping Detection. Curr Sports Med Rep. Apr;18:97–104. Epub 2019/04/11.

Thevis M, Sauer M, Geyer H, Sigmund G, Mareck U, Schanzer W. 2008. Determination of the prevalence of anabolic steroids, stimulants, and selected drugs subject to doping controls among elite sport students using analytical chemistry. J Sports Sci. Aug;26:1059–1065.

Ulrich R, Pope HG, Jr., Cleret L, Petroczi A, Nepusz T, Schaffer J, Kanayama G, Comstock RD, Simon P. 2018. Doping in Two Elite Athletics Competitions Assessed by Randomized-Response Surveys. Sports Med. Jan;48:211–219.

WADA. 2009. Blood Analytical Requirements for the Athlete Biological Passport TD2010BAR 1.0. 1.06.2019;Version 1.0.

2017 Anti-Doping test figures. 23.08.2018 ed. Montreal: WADA. Available from https://www.wada-ama.org/en/resources/laboratories/anti-doping-testing-figures

